# Endogenous labeling empowers accurate detection of m^6^A from single long reads of direct RNA sequencing

**DOI:** 10.1101/2024.01.30.577990

**Authors:** Wenbing Guo, Zhijun Ren, Xiang Huang, Jialiang He, Jie Zhang, Zehong Wu, Yang Guo, Zijun Zhang, Yixian Cun, Jinkai Wang

**Author notes:** These authors contributed equally (Wenbing Guo, Zhijun Ren, Xiang Huang). Corresponding author.; Mailing address: No. 74, Zhongshan Rd.2, Guangzhou 510080, China.

## Abstract

Although plenty of machine learning models have been developed to detect m^6^A RNA modification sites using the electric current signals of ONT direct RNA sequencing (DRS) reads, the landscape of m^6^A on different RNA isoforms is still a mystery due to their limited capacity to distinguish the m^6^A on individual long reads and RNA isoforms. The primary challenge in training the model with single-read accuracy is the difficulty of obtaining the training data from individual DRS reads that comprehensively represent the m^6^A on endogenous RNAs. Here, we endogenously label the methylated m^6^A sites on single ONT DRS reads by APOBEC1-YTH induced C-to-U mutations, strategically positioned 10-100 nt away from the known m^6^A sites on the same reads. Adopting a semi-supervised leaning strategy, we obtain 700,438 reliable 5-mer single-read level m^6^A signals, providing a comprehensive representation of m^6^A on endogenous RNAs. Leveraging this dataset, we develop m6Aiso, a deep residual neural network model that not only accurately identifies and quantifies known m^6^A sites but also reveals unknown, subtly methylated m^6^A sites responsive to METTL3 depletion. Analyzing m6Aiso-determined m^6^A on single reads and isoforms uncovers distance-dependent linkages of m^6^A sites along single molecules, as well as differential methylation of identical m^6^A sites on different isoforms. Moreover, we find wide-spread functionally important dynamic changes of m^6^A sites on specific isoforms during epithelial-mesenchymal transition (EMT). The pivotal utilization of the endogenous labeling strategy empowers m6Aiso to achieve remarkable precision in pinpointing m^6^A on individual molecules, underscores its effectiveness in elucidating the intricate dynamics and complexities of m^6^A across RNA isoforms.

## INTRODUCTION

N^6^-methyadenosine (m^6^A) is a prevalent and dynamic modification on mRNAs and diverse types of noncoding RNAs^1^. It is mainly catalyzed by the m^6^A methyltransferase METTL3 at the DRACH (D = A, G or U; R = A or G; H = A, C or U) motif on RNAs^2^. Various m^6^A reader proteins, notably the YTH family proteins, play important roles in governing a wide range of cellular processes through reading m^6^A, including degradation, translation, transcription, polyadenylation, nuclear export, splicing, and et. al.^3–7^ In recent years, we have gained comprehensive understanding of the functions and distributions of m^6^A sites due to the rapid development of the technologies that identify the locations of the m^6^A sites^8–11^. Particularly, a recently developed method GLORI has been proved to be able to provide accurate identification and absolute quantification of m^6^A at single-nucleotide resolution through glyoxal and nitrite-mediated deamination of unmethylated adenosine^10^. However, the majority of these methods suffer from the ambiguity of m^6^A modifications on isoforms due to their inability to preserve intact RNAs. Furthermore, more and more researches emphasize the intricate association between m^6^A and the generation and metabolism of RNA isoforms^12–14^. Especially, recent studies have revealed that the exon junction complex (EJC) plays an inhibitory role in the deposition of m^6^A near splicing junctions, resulting in significant depletion of m^6^A sites in exon regions within 200 nt of internal exon boundaries and overrepresentation of m^6^A sites in long internal exons^15–17^. These findings strongly suggest the possibility of selective m^6^A deposition on different mRNA isoforms even at the exactly same locations. However, due to the lack of methods to accurately identify the m^6^A on each intact RNA molecule, how the m^6^A sites are distributed among various RNA isoforms remains unknown.

The third generation long-read sequencing technology has been proven to be a powerful strategy for deciphering the complexities of RNA isoforms^18, 19^, thus detecting m^6^A RNA modifications on the long-reads can provide the m^6^A locations in the context of RNA isoforms. Particularly, the Oxford Nanopore (ONT) direct RNA sequencing (DRS)^20^ captures the electric current changes for each continuous five nucleotides on RNAs when they traverse the nanopore. A variety of machine learning models have been developed with the aim to identify the location of single-nucleotide m^6^A sites on the genome without information on single reads. These strategies can be broadly grouped into two categories based on the training labels. The first category relies on *in vitro* transcripts (IVTs), where all “A” residues are substituted with m^6^A residues using N^6^-methyladenosine-5′ triphosphate(m^6^ATP) instead of ATP through *in vitro* transcription, such as Epinano^21^, ELIGOS^22^, Nanocompore^23^, and TandemMod^24^. The second category of models take advantage of the distribution of current signals of all reads covering the annotated single-nucleotide m^6^A sites in cells, such as xPore^25^ and MINES^26^.

In despite of the rapid development of the machine learning algorithms that predict m^6^A sites using ONT DRS reads, training a model to accurately identify m^6^A at single DRS reads is still of great challenge due to the lack of high-quality m^6^A modified DRS signals on single molecules. There are several different strategies to obtain single-molecule m^6^A signals. Nanom6A^27^ leverages IVT RNAs and was trained using DRS data that only includes 130 sites of the RRACH motif. These sites have limited diversity in the sequence context, and it methylates all adenosines without selectivity, thus causing superimposed effect on signal readout for the m^6^A motifs with multiple adenosines^28^. Another method, DENA^28^, is trained by comparing base calling errors in individual reads obtained from wild-type (WT) and m^6^A methyltransferase knockout (KO) conditions in *Arabidopsis thaliana* across various RRACH motifs. This strategy has to suffer from the low accuracy of the labels. The most recently published method m6Anet^29^ used a multiple instance learning framework on the DRS reads covering the highly methylated m^6^A sites revealed by m6ACE-seq. It outperforms all other published methods in terms of predicting highly comparable m^6^A sites with the experimental methods, as demonstrated by a third-party evaluation^30^. However, the single-read accuracy of these methods has to suffer from the intrinsically biased representation of the available m^6^A signals. Therefore, there is an urgent need for comprehensive and unbiased labeling of m^6^A on individual cellular RNA molecules to ultimately train a model with reliable single-read accuracy.

DART-seq, which was reported to induce C-to-U mutations right beside the m^6^A sites through APOBEC1-YTH fusion protein in live cells, was previously recognized to be a method that identifies m^6^A at single-nucleotide resolution^31^. However, although the C-to-U mutations can label the m^6^A endogenously on the same molecules, the C-to-U mutations beside the m^6^A sites must distort the signals of m^6^A modified 5-mers, which constitute the smallest unit for ONT signal. In this study, we revealed that APOBEC1-YTH is also possible to induce clustered C-to-U mutations away from the m^6^A sites without distorting the 5-mers that cover the GLORI identified m^6^A sites on the same molecules. Based on a cautiously verified strategy to determine the methylation status of GLORI annotated m^6^A sites from the C-to-U mutations on the same DRS reads, we finally obtained 700,438 undistorted m^6^A signals on single reads at the GLORI annotated m^6^A sites. Through convolutional neural networks (CNN) deep learning based on these DRS m^6^A signals, we developed m^6^Aiso, which accurately identified m^6^A on single reads and revealed complexity of m^6^A on RNA isoforms as well as the functional relevant isoform-specific m^6^A dynamics during Epithelial-mesenchymal transition (EMT).

## RESULTS

### Determination of positive m^6^A signals on single DRS reads through endogenous labelling

Upon reanalyzing the data of original DART-seq^31^, we realized that although 59.7% of C-to-U mutations were not within DRACH m^6^A motifs (Supplementary Fig. 1a), 39.5% of which were actually located within 100 nt of the GLORI identified m^6^A sites in the same cell line (Supplementary Fig. 1b, c), suggesting that APOBEC1-YTH induces C-to-U mutations mostly within 100 nt of m^6^A sites. Therefore, these C-to-U mutations away from the m^6^A sites may be used to label the m^6^A without interfering with the electric current of m^6^A methylated 5-mers.

To efficiently label the m^6^A endogenously on single RNAs, we performed ONT direct RNA sequencing deeply on the mRNAs of HEK293T cells with efficient translation of copGFP-tagged APOBEC1-YTH (5,503,279 reads) and copGFP-tagged empty vector (4,932,654 reads). After filtered out the known SNPs as well as the mutations that could be induced by APOBEC1 alone^31^ or those mutations occurred in empty vector, we identified 384,096 C-to-U mutations from DRS long reads. These mutations were distributed in a manner resembling the distribution of m^6^A sites with an enrichment near stop codons (Supplementary Fig. 1d), although exhibiting a slight 3’ bias, which is consistent with the general bias observed in ONT reads^32^. We found 166,507 (43.4%) of the C-to-U mutations, including those not within m^6^A motifs, were clustered within 100 nt. Furthermore, the clustered rather than the non-clustered C-to-U mutations were enriched near the stop codons of mRNAs (Supplementary Fig. 1e). These results indicate that APOBEC1-YTH tends to induce clustered C-to-U mutations other than singleton mutations around the genuine m^6^A sites.

Thereby, we proceeded to determine the proper single-read m^6^A signals at the GLORI identified m^6^A sites by requiring the occurrence of at least one clustered C-to-U mutation 10-100 nt away, but no mutation occurring less than 10 nt away from the m^6^A sites on the same reads (Fig. 1a). By assuming that the C-to-U mutations were due to the m^6^A methylation of the nearest m^6^A sites on the same RNAs, we determined 1,633,238 m^6^A signals on 1,787,651 single DRS reads at 15,534 unique GLORI identified m^6^A sites in HEK293T. Meanwhile, from the same DRS dataset, we also identified 18,630,201 single-read level unmodified signals at the DRACH motifs no less than 20 nt away from any m^6^A sites annotated by miCLIP^33^, GLORI^10^, m6A-SAC-seq^9^, and m6ACE-seq^34^, nor within any m^6^A peaks of m6A-seq^35^ in HEK293T cells.

**Figure 1.**
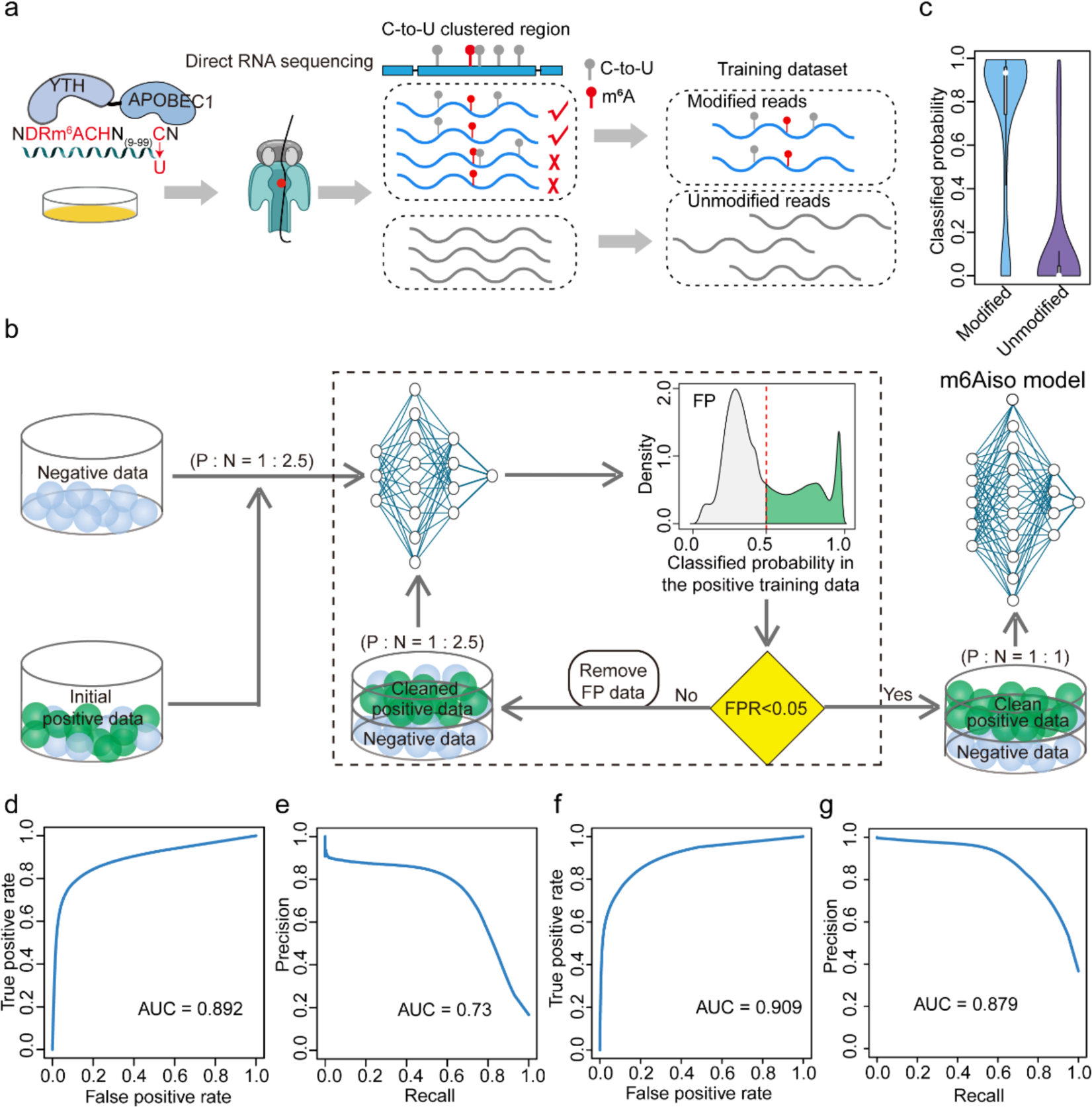
Endogenously labeling of m^6^A on single reads and assessment of m6Aiso at single molecule level. **a**, Flowcharts illustrating the classification of modified and unmodified native reads. **b**, Schematic illustrating a semi-supervised leaning strategy for filtering out false positive signals. P:N, positive data: negative data; FP, false positive; FPR, false positive rate. **c**, The distribution of predicted modification probabilities using the m6Aiso model in the independent validation dataset. **d, e**, Receiver operating characteristic (ROC) curve (**d**) and precision-recall (PR) curve (**e**) of m6Aiso in the independent validation dataset. **f**, **g**, ROC curve (**f**) and PR curve (**g**) of m6Aiso single-molecule prediction on synthesized RNA reads from the Curlcake datasets.

Based on the m^6^A modified and unmodified signals at single reads, we developed a CNN-based model named m6Aiso to predict m^6^A modifications at the single molecular level (Supplementary Fig. 2). Because the correct m^6^A sites responsible for the C-to-U mutations were sometimes ambiguous especially when multiple GLORI identified m^6^A sites were near the mutations, we recognized that there should be some false positive signals in the positive training data. Being aware of the existence of these false positives, we used a semi-supervised leaning strategy to train a deep residual neural network model (ResNet) with the input of 7 nt RNA sequences (from 1 nt upstream to 1 nt downstream of DRACH) and their local electric current signal features to predict the methylation states of the DRACH motifs on single reads (Fig. 1b). In this training process, an initial model of m6Aiso was trained using all the training data. We then discarded the samples with modified probabilities < 0.5 from the positive data set and updated the training data to re-train the m6Aiso model. This update and re-training processes were repeated and terminated till the false positive rate (FPR) < 0.05 in the positive training data. We ultimately preserved 700,438 (42.9%) clean single-read m^6^A modified signals at 13,954 unique m^6^A sites for the final training of m6Aiso. Although not gold standard, we still found 77.1% of these clean m^6^A signals were also supported by the previous published m6Anet^29^, suggesting the high quality of these signals.

These 13,954 unique m^6^A sites that derived from the clean positive single-read m^6^A signals had the similar compositions of m^6^A motifs as well as distributions of m^6^A level with GLORI identified m^6^A sites in HEK293T cells^10^ (Supplementary Fig. 1f, g). Of note, 126,030 (18.0%) of these single-read signals were from lowly methylated m^6^A sites (level < 0.25), which were largely omitted in the training data of previous methods (Supplementary Fig. 1h). In conclusion, our positive training dataset integrally represents the endogenous m^6^A sites with various stoichiometry and sequence contexts. Therefore, the model trained using these data should be less prone to unwarranted biases, such as the difficulty in training for lowly modified m^6^A sites.

To evaluate the performance of m6Aiso, we used an independent flow cell of ONT DRS reads obtained from HEK293T cells infected with empty vector. The validation dataset comprises all the 171,791 single-read signals at 3,265 GLORI determined extremely highly methylated (level > 0.95) m^6^A sites in HEK293T cells as positive data as well as 858,955 nonmodified single-read signals as the negative data. The m6Aiso determined modification probabilities were concentrated at the expected extremes in both unmodified and modified reads in the validation dataset (Fig. 1c). Based on this validation dataset composed of only signals from extremely highly methylated m^6^A sites, m6Aiso achieved a receiver operating characteristic (ROC) AUC of 0.892 and a precision-recall (PR) AUC of 0.73 (Fig. 1d, e). Moreover, m6Aiso achieved an even higher ROC AUC of 0.909 and a PR AUC of 0.879 (Fig. 1f, g) based on the synthetic IVT data with four types of valid m^6^A motifs^21^.

### m6Aiso accurately detects and quantifies both highly and lowly methylated m^6^A sites

By using m6Aiso, we then identified 35,074 m^6^A sites with strong enrichment near stop codons in HEK293T cells infected with empty vector (Fig. 2a). These m6Aiso identified m^6^A sites resembled the 5-mers profile of GLORI identified m^6^A sites (Fig. 2b); moreover, in contrast to m6Anet^29^, which appears to mostly capture the m^6^A sites with levels greater than 0.4, m6Aiso identified m^6^A sites in HEK293T cells showed a ‘saddle-shaped’ distribution of m^6^A levels with 15,469 (44.1%) of them were under 0.25, which is highly consistent with GLORI determined m^6^A levels (Fig. 2c).

**Figure 2.**
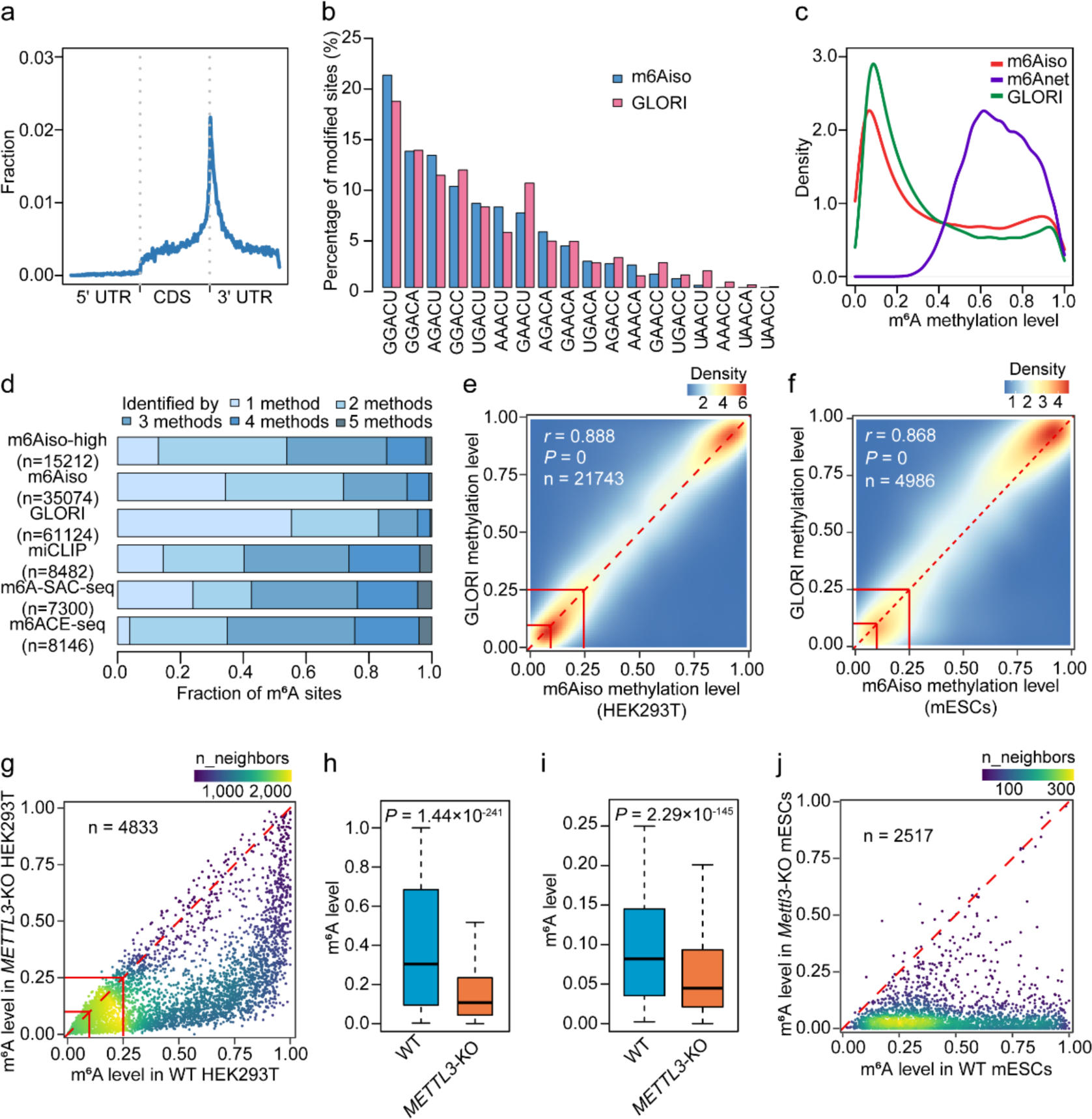
m6Aiso accurately detects and quantifies both highly and lowly methylated m^6^A sites. **a**, Metagene profile illustrating the distribution of the modified sites captured by m6Aiso. **b**, Comparison of the percentages of modified sites predicted as modified by m6Aiso and by GLORI on the DRACH motifs. **c**, Density plot comparing the distributions of m^6^A methylation levels determined by GLORI (green line), m6Aiso (red line), and m6Anet (purple line). **d**, Percentages for m^6^A sites detected by different number of methods, including m6Aiso as well as GLORI, miCLIP, m6ACE-seq and m6A-SAC-seq. “m6Aiso-high” denotes the m6Aiso determined m^6^A sites with methylation levels > 0.4. **e**, Correlation of the m^6^A levels estimated by m6Aiso and GLORI in HEK293T cells infected with empty vector. The m^6^A levels of 0.1 and 0.25 are indicated by red lines. **f**, Correlation of the m^6^A levels estimated by m6Aiso and GLORI in mESCs. **g**, **h**, Scatter plot (**g**) and box plot (**h**) demonstrating a remarkable reduction of methylation levels in *METTL3*-KO HEK293T cells compared to WT HEK293T cells for all the m6Aiso determined m^6^A sites. *P* value was calculated by two-tailed Wilcoxon rank-sum test. **i**, Box plot illustrating a decrease in methylation levels in *METTL3*-KO HEK293T cells compared to WT HEK293T cells for the m6Aiso determined lowly methylated m^6^A sites (level < 0.25). *P* value was calculated by two-tailed Wilcoxon rank-sum test. **j**, Scatter plot depicting a remarkable reduction of methylation levels in *Mettl3*-KO mESCs compared to WT mESCs for the GLORI-annotated m^6^A sites. For boxplots: center line, median; box limits, upper and lower quartiles; whiskers, 1.5x interquartile range.

To further evaluate the accuracy of m6Aiso identified m^6^A sites, we took advantage of the previously known m^6^A sites in HEK293T cells identified by GLORI^10^, miCLIP^33^, m6A-SAC-seq^9^, and m6ACE-seq^34^, which are the experimental methods that can accurately identify m^6^A in single-nucleotide resolution. We found 23,044 (65.7%) of m6Aiso identified m^6^A sites could be validated by at least one of the four experimental methods, while 12,030 (34.3%) m^6^A sites are m6Aiso-specific (Fig. 2d). In contrast to the shared m^6^A sites, we found these m6Aiso-specific sites showed very low m^6^A stoichiometry with 74.6% of them have m^6^A levels under 0.25 (Supplementary Fig. 3a). In addition, if we only consider the 15,212 sites with m^6^A levels greater than 0.4 to mimic the level distribution of m6Anet identified m^6^A sites, 82.0% of m^6^Aiso determined m^6^A sites were GLORI identified m^6^A sites in HEK293T, which was comparable with m6Anet but much higher than DENA and nanom6A (Supplementary Fig. 3b). On the other hand, although methylated lowly, these 12,030 m6Aiso-specific sites still displayed a strong enrichment near the stop codons (Supplementary Fig. 3c), and 5,777 (48.0%) were annotated within m6A-Atlas (v2.0) database^36^, which contains the known m^6^A sites in diverse cell lines and tissues. These results suggest that the m6Aiso-specific m^6^A predictions still represent the authentic lowly methylated m^6^A sites that were technically difficult to be confidently determined by previous methods.

We then tested how accurate was m6Aiso in quantifying the m^6^A levels by comparing with the m^6^A levels measured by GLORI in HEK293T cells and mouse embryonic stem cells (mESCs). We found a strong correlation of m6Aiso determined m^6^A levels with GLORI determined m^6^A levels in HEK293T (Pearson’ *r* = 0.888, Fig. 2e), outperforming m6Anet, DENA and nanom6A (Supplementary Fig. 3d-f). Of note, m^6^A sites with levels under 0.25 were also highly consistent with GLORI, suggesting m6Aiso also has sufficient power and accuracy to quantify the lowly methylated m^6^A sites. Similar results were also observed in mESCs (Pearson’ *r* = 0.868, Fig. 3f). When we classified the m^6^A sites by the 5-mers, we found the correlation of m^6^A levels between m6Aiso and GLORI in HEK293T cells was particularly robust for the common motifs (Supplementary Fig. 4) (for example, GGACU, *r* = 0.944; GGACC, *r* = 0.937; AGACU, *r* = 0.931) and outperformed m6Anet^29^ as well as mAFiA^37^, a model trained with synthetic RNAs comprised of 6 different 5-mers of m^6^A motif and reported on bioRxiv.

**Figure 3.**
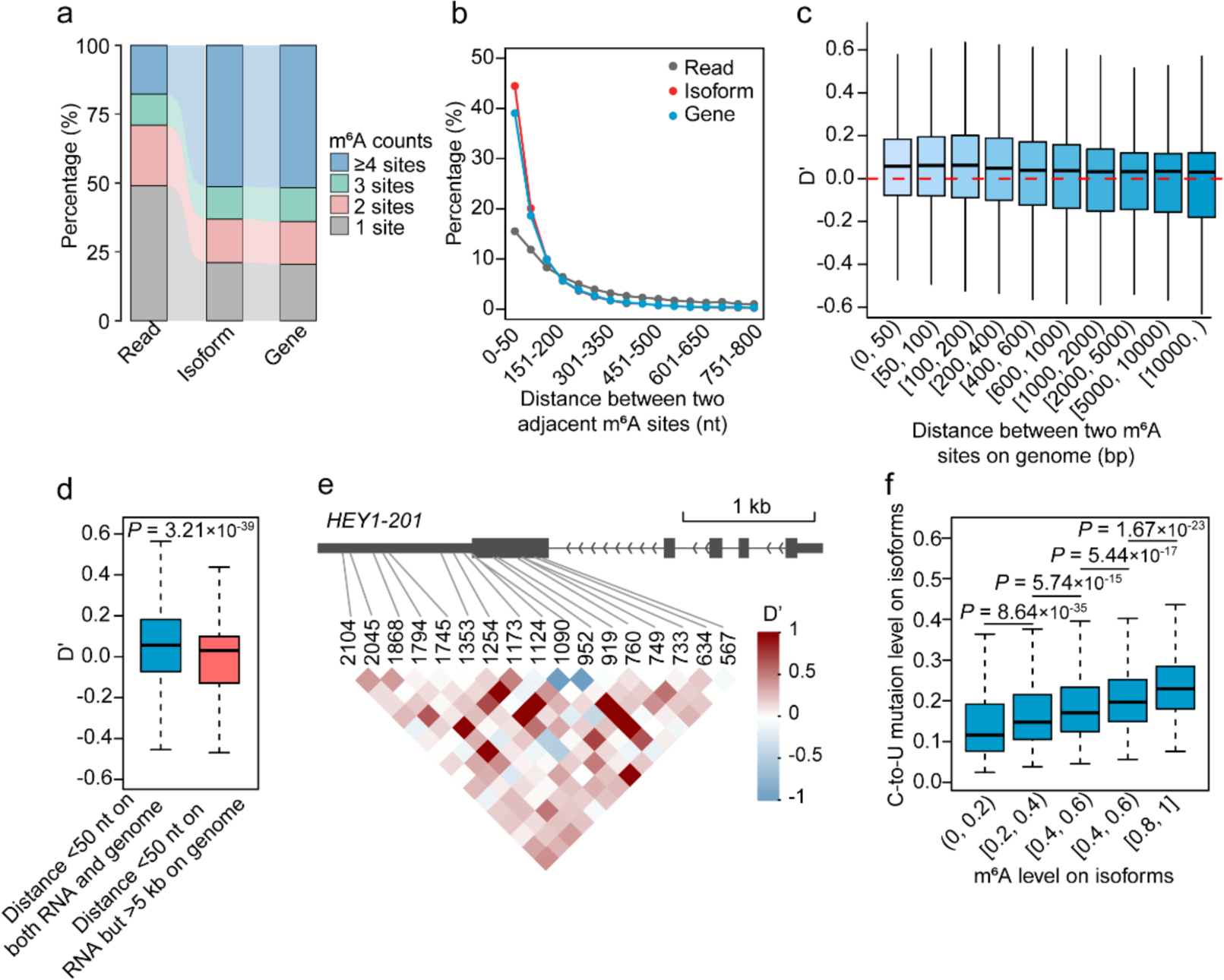
Linkage of m^6^A sites along the same RNA molecules. **a**, Sankey diagram comparing the percentages of single reads, isoforms, and genes with different number of m^6^A sites. **b**, Line chart showing the percentages of m^6^A sites within various distances to adjacent m^6^A sites at the read (gray line), isoform (red line) and gene (blue line) level. **c**, Box plot showing the linkage (D’) of pairs of m^6^A sites with different distances on the genome. **d**, Box plot comparing the D’ for pairs of m^6^A sites with distance < 50 nt on RNAs but > 5 kb on genes as compared with those with distance < 50 nt on both RNAs and genes. *P* value was calculated by two-tailed Wilcoxon rank-sum test. **e**, Representative example of the coordinated occurrence of m^6^A sites in *HEY1-201* transcript. **f**, Correlation of isoform m^6^A levels determined by m6Aiso and APOBEC1-YTH induced C-to-U mutations. *P* values were calculated by two-tailed Wilcoxon test. For boxplots: center line, median; box limits, upper and lower quartiles; whiskers, 1.5x interquartile range.

We next evaluated the accuracy of m6Aiso by using the previously published ONT DRS data of *METTL3*-KO and wild-type (WT) in HEK293T cells^38^ and mESCs^30^, respectively. First of all, we found highly consistent of m6Aiso determined m^6^A levels (Pearson’ *r* = 0.983, Supplementary Fig. 3g) between the two replicates of WT HEK293T. We then observed robust downregulation of m6Aiso determined m^6^A levels in *METTL3*-KO for the vast majority of m^6^A sites (Fig. 2g, h). Of note, we found the m^6^A downregulation were more pronounced as determined by m6Aiso than m6Anet (Supplementary Fig. 3h, i). Moreover, the majority of the above described m6Aiso-specific m6A sites and lowly methylated m^6^A sites (level < 0.25) were also down-regulated, including those with levels < 0.1 (Fig. 2i). Similar results were also observed for *Mettl3* KO in mESCs (Fig. 2j and Supplementary Fig. 3j-m). These results suggest that m6Aiso not only accurately detects the known m^6^A sites but also the genuine lowly methylated m^6^A sites that are largely overlooked by previous methods.

### Genomic distance dependent linkage of m^6^A sites along single molecules

It has been reported that a significant portion of m^6^A sites tend to be clustered in short regions^9, 10, 33^. However, it remains unclear whether the m^6^A sites are clustered on isoforms and single molecules. By using m6Aiso, we identified a total of 35,074 m^6^A sites on 13,755 RNA isoforms of 6,668 genes in HEK293T cells transfected with empty vector. There were on average 5.1, 5.1 and 2.3 m^6^A sites for each gene, isoform, and read, respectively (Supplementary Fig. 5a-c); of note, in contract to the genes and isoforms with <25% of them had single methylated m^6^A sites, 49.0% of methylated DRS reads had single methylated m^6^A sites (Fig. 3a). Moreover, consistent with the previous reports^9, 10, 33^, 39.1% of m^6^A modifications were clustered with adjacent m^6^A sites within 50 bp on the same genes, and it was 44.5% on the same isoforms (Fig. 3b). However, only 15.5% of the m^6^A sites were clustered within 50 bp on the same ONT reads, suggesting that the m^6^A sites are not likely to be strongly linked on the same RNA molecules (Fig. 3b).

We then employed the parameters D’, a matric commonly used to assess the linkage disequilibrium (LD) between two genetic variants within a population^39^, to test whether the different m^6^A sites of the same RNA isoforms tend to be linked on the same reads or simply cooccur on the same reads due to random chances. As shown in Figure 3c, we observed a relatively weak but nonnegligible linkages between m^6^A sites with genomic distances less than 200 bp, but the linkage decreased gradually with the distances increases and it became not very recognizable when the distances were greater than 1 kb. Consistent with the previous report that m^6^A is deposited co-transcriptionally on chromatin RNAs^40^, we found the pairs of m^6^A sites with distances < 50 bp on RNAs but > 5 kb on genome had significantly smaller D’ than the pairs of sites with distances < 50 bp on both RNAs and genome (Fig. 3d). As exemplified in Figure 3e, 17 m^6^A sites in the last exon of *HEY1-201* isoform are linked as a block with strong D’ for certain pairs of m^6^A sites, suggesting the existence of strongly linked m^6^A sites although in general the linkage is relatively weak across the transcriptome. Therefore, the m^6^A sites are partially linked along single RNA molecules in a manner that depends on the distances on genome other than on RNAs.

### m6Aiso reveals differential methylation of identical m^6^A sites on different isoforms

To test whether m6Aiso can accurately quantity m^6^A on each RNA isoform, we calculated the m^6^A levels of each isoform as the mean level of all the m^6^A sites. We found the average mutation rates of APOBEC-YTH induced C-to-U mutations of individual isoforms had a significant positive correlation with the m6Aiso other than m6Anet determined m^6^A levels of isoforms in HEK293T cells (Fig. 3f and Supplementary Fig. 5d). Moreover, the m6Aiso determined isoform m^6^A levels had significant negative correlation with the expression of isoforms (Supplementary Fig. 5e), which was consistent with the well-characterized function of m^6^A in promoting RNA decay^41, 42^. In addition, consistent with the previous reports that EJCs suppress the m^6^A deposition within 200 bp of exon boundaries^15–17^, we also observed lowly methylation of m^6^A within 200 bp of exon boundaries on the isoforms (Fig. 4a) We were then interested in whether the same m^6^A sties with the identical genomic locations can also be methylated differentially on different isoforms. By comparing the m^6^A levels of the same m^6^A sites among different RNA isoforms, we determined 716 (unique, n = 187) and 626 (unique, n = 89) m^6^A sites that were methylated significantly higher and lower in specific isoforms than the combination of other isoforms of the same genes (Fig. 4b). As compared with the isoforms with isoform-specific lowly methylated m^6^A sites, the isoforms with isoform-specific highly methylated m^6^A sites were significantly enriched in those isoforms with retained introns (*P* = 0.0131, Fig. 4c). Furthermore, we observed the m^6^A sites on intron-retained isoforms were tended to located farther away from the exon junctions than other transcripts (Supplementary Fig. 5f). To test whether the inhibitory role of EJC on m^6^A results in differential methylation of m^6^A sites on the alternatively spliced isoforms, we determined the m^6^A sites with alternative exon junction distances (EJDs) on different isoforms. As compared with the levels of these m^6^A sites on isoforms with EJD < 100 bp, we found 30 and 2 unique m^6^A sites were methylated higher and lower on isoforms with EJD > 200, respectively (Fig. 4d). These results suggest the inhibitory role of EJC can account for at least a part of isoform-specific m^6^A methylation. As exemplified in Figure 4e, two m^6^A in *SMUG1* were m^6^A methylated higher on the isoform with a retained intron (*SMUG1-215*) than the other isoform with the intron spliced (*SMUG1-201*) (Fig. 4 e, f).

**Figure 4.**
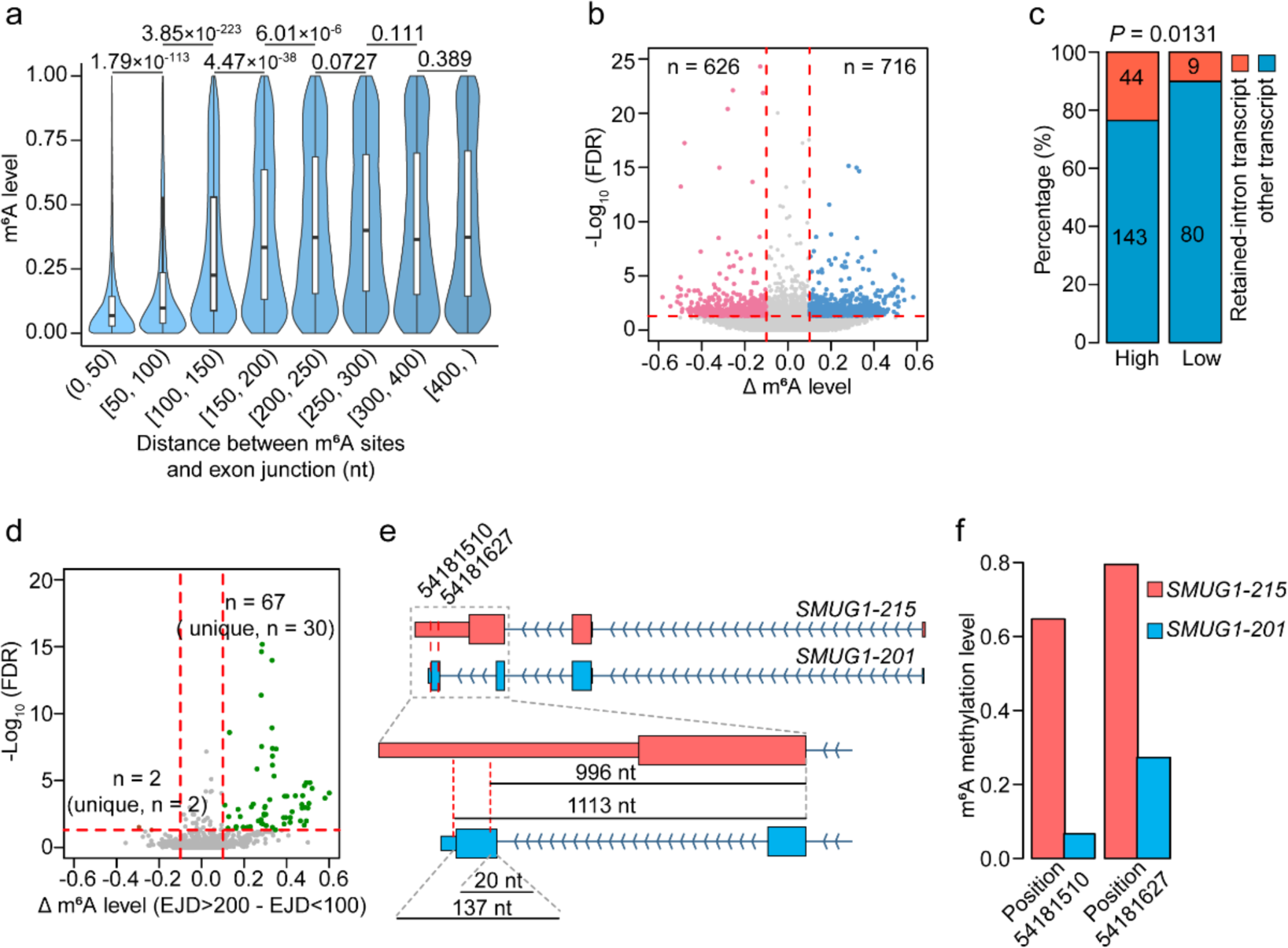
m6Aiso identified m^6^A sites on different isoforms of the same genes. **a**, Violin plot depicting the correlation between methylation levels of m^6^A sites and their proximity to the exon junctions. **b**, Volcano plot depicting the differential methylation of single-nucleotide m^6^A sites on specific isoforms as compared with the m^6^A level of the identical sites on the combination of all other isoforms in the same genes. **c**, Bar plot depicting the number of retained-intron transcripts associated with isoform-specific lowly methylated and highly methylated m^6^A sites. *P* value was calculated by two-tailed Chi-square test. **d**, Volcano plot showing the differential m^6^A methylation at identical m^6^A sites on different isoforms with exon-junction distances > 200 nt and < 100 nt. **e**, **f**, Location diagram (**e**) and box plot (**f**) showing the differential m^6^A methylation at identical m^6^A sites on two isoforms with varying distances from exon junction due to intron retention.

### m6Aiso reveals isoform-specific dynamic changes of m^6^A during epithelial-mesenchymal transition

Recent studies have unveiled the pivotal roles of m^6^A in the process of epithelial-mesenchymal transition (EMT)^43, 44^, which is also a process critically regulated by alternative RNA splicing^45^. To investigate whether m^6^A can be dynamically regulated in an isoform-specific manner, we performed ONT DRS to examine the m^6^A changes on specific RNA isoforms in HeLa cells upon TGF-β induced EMT. First of all, we confirmed the EMT was successfully induced through western blot and RNA-seq based expression analysis (Supplementary Fig. 6a-c). Through identification of the m^6^A modified adenosines on each single DRS reads using m6Aiso, we determined 34,418 and 33,982 unique m^6^A sites in the control and TGF-β samples, respectively (Fig. 5a). These m^6^A sites were strongly enriched near stop codons with 84% of the sites were overlapped between control and TGF-β samples (Fig. 5a, b). Furthermore, we observed a significant global increase of m^6^A levels in TGF-β samples (*P* = 1.21×10^-13^, Fig. 5c). Consistently, we determined 1,013 up-regulated m^6^A sites on 906 unique RNA isoforms and 208 down-regulated m^6^A sites on 207 RNA isoforms upon TGF-β induction of EMT (Fig. 5d, e). The genes with up-regulated m^6^A on isoforms showed significant enrichment for EMT related Gene Ontology (GO) terms, such as “Cellular response to transforming growth factor beta stimulus” and “Beta-catenin binding” (Fig. 5f).

**Figure 5.**
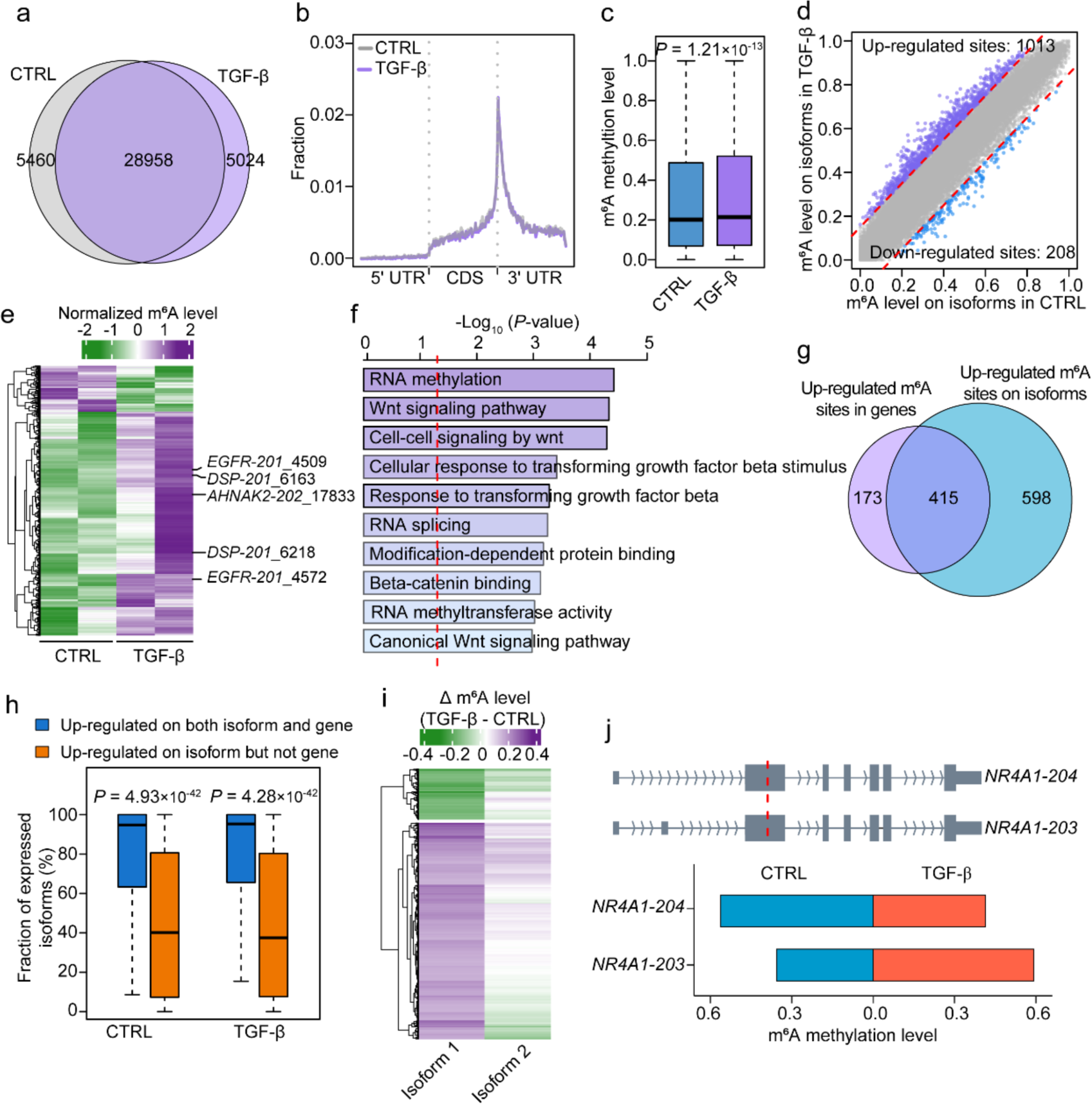
Dynamical regulation of m^6^A in an isoform-specific manner during TGF-β induced EMT. **a**, Venn diagram comparing the number of m^6^A sites detected by m6Aiso in control and TGF-β induced cells. CTRL, control. **b**, Metagene profiles depicting the distributions of m^6^A sites in control (gray line) and TGF-β induced cells (purple line). **c**, Box plot comparing the m^6^A levels between control and TGF-β induced cells. **d**, Scatter plot depicting the differential m^6^A sites on isoforms between control and TGF-β induced cells. Purple and blue dots denote up-regulated and down-regulated m^6^A sites during EMT process, respectively. **e**, Heatmap showing the methylation levels of differentially methylated m^6^A sites between control and TGF-β induced cells. **f**, GO enrichment for the isoforms with up-regulated m^6^A sites. **g**, Venn diagram comparing the up-regulated m^6^A sites at the gene and isoform levels. **h**, Box plot comparing the expression proportions of specific isoforms out of all isoforms of the same genes for the isoforms with up-regulated m^6^A sites at both the isoform and gene levels with the isoforms with up-regulated m^6^A sites at the isoform level but not the gene level. *P* values were calculated by two-tailed Wilcoxon rank-sum test. **i**, Heatmap comparing the changes of methylation levels of identical m^6^A sites on two different isoforms upon TGF-β induction. Isoform 1 denotes the isoforms with differentially methylated m^6^A sites, while isoform 2 denotes the randomly selected other isoforms originated from the same genes as isoform 1. **j**, Representative example of an m^6^A site in *NR4A1* that changes to different directions on two different isoforms during TGF-β induction. For boxplots: center line, median; box limits, upper and lower quartiles; whiskers, 1.5x interquartile range.

To test whether the m^6^A changes in EMT are isoform-specific, we compared the differential m^6^A sites determined at gene and isoform levels. Of note, we found 598 (59%) of the 1,013 up-regulated sites on isoforms were overlooked at the gene level comparison (Fig. 5g). In contrast to the shared differential m^6^A sites on both gene and isoform levels, the differentially methylated sites that exclusively determined on isoform level were strongly enriched in those minor isoforms that expressed lower than the major isoforms of the same genes, suggesting the isoform-based m^6^A analyses are critical in deciphering the dynamics of m^6^A on minor isoforms (Fig. 5h). Similarly, 158 (76%) of the 208 down-regulated sites on isoforms were also overlooked at the gene level (Supplementary Fig. 6f). For the m^6^A sites that were only differentially methylated on certain RNA isoforms other than genes, we found the randomly selected RNA isoforms of the same genes did not show significant m^6^A changes at the same m^6^A sites (Fig. 5i), suggesting that these m^6^A sites are differentially methylated upon TGF-β induction in an isoform-specific manner. As exemplified in Figure 5j, an m^6^A site in the long internal exon of *NR4A1*, an important enhancer for TGF-βinduced EMT^46^, were down-regulated in the isoform *NR4A1-204* but up-regulated in the isoform *NR4A1-203*, suggesting isoform-specific dynamic changes of m^6^A in EMT.

## DISCUSSION

In this study, we developed an endogenous labeling-based method to generate a large amount of endogenous m^6^A signals of ONT DRS reads. We then trained a CNN-based deep leaning model named m6Aiso to accurately identify m^6^A at single reads of ONT and revealed the diversity of m^6^A on different RNA isoforms. In contrast to previous ONT DRS-based methods, m6Aiso has obtained exceedingly accuracy of detecting m^6^A on single reads and is able to distinguish the m^6^A deposited on different RNA isoforms.

Lowly methylated m^6^A sites extremely important in decoding the m^6^A on RNA isoforms, especially those extensively existed minor isoforms. Previous methods including experimental and ONT DRS-based methods do not have sufficient power to distinguish the lowly methylated sites from noises. Even GLORI has to remove all the sites with levels < 0.1 to avoid the inclusion of extensive false positives due to incomplete A-to-I conversion of unmodified adenosines^10^. ONT DRS-based methods, such as m6Anet^29^, a model trained from the single-read m^6^A signals from the highly methylated sites, omitted most of the m^6^A sites with m^6^A levels < 0.4. Surprisingly, we found m6Aiso exhibited reliable ability of detecting the lowly methylated m^6^A sites. Although a large fraction of the m6Aiso determined lowly methylated m^6^A cannot be identified by GLORI, these m^6^A sites are still distributed as genuine m^6^A sites and are mostly sensitive to METTL3 depletion, including the m^6^A sites with levels < 0.1. On the other hand, highly and lowly methylated m^6^A sites may have intrinsic differences of m^6^A signals. The differences would be even significant between the m^6^A signals on endogenous RNAs and synthetic RNAs, which usually have limited types of motifs and less complexity of sequences. Indeed, previous studies have reported that the stoichiometry of m^6^A sites can be predicted by the surrounding sequences alone^47, 48^, suggesting the machine learning model might take advantage of the sequences or other unexpected features in predicting m^6^A especially for the sites that are insufficiently represented in the training data. Therefore, a large amount of m^6^A signals on single reads with comprehensive representation of endogenous m^6^A sites and stoichiometries are critical in training a highly accurate and powerful model in detecting m^6^A using ONT DRS reads. Fortunately, our training data are comprised of 700,438 positive single-read m^6^A signals on the m^6^A sites with similar distribution of motifs and levels as the endogenous m^6^A sites. This is mostly because the APOBEC1-YTH targets the individual RNA molecules independently, thus it provides independent positive m^6^A signals on individual reads in a way like randomly samplings of the endogenous m^6^A modified RNAs.

In this study, we revealed the existence of isoform-specific m^6^A modification and dynamic changes even at the identical m^6^A sites. This finding highlights the benefits of using ONT DRS to identify the locations and dynamics of m^6^A, especially when we have developed m6Aiso to identify the m^6^A with high accuracy and power. We found a part of isoform-specific m^6^A sites are due to alternative exon junction distances from the same m^6^A sites on different RNA isoforms generated by alternative splicing, suggesting that alternative splicing can regulate alternative m^6^A deposition on different isoforms through EJC. On the other hand, only a part of the alternatively methylated m^6^A sites have alternative exon junction distances, suggesting that EJC only accounts for a small part of alternatively methylated m^6^A sites. This observation is highly consistent with a recent study reporting that EJC only partially contributes to the inhibition of m^6^A at exon-intron boundaries for some short internal exons^49^.

Therefore, there should also be additional mechanisms that can mediate the isoform-specific m^6^A deposition. Indeed, many studies have reported the role of m^6^A in regulating alternative splicing^4, 7, 12, 50^. On the other hand, previous studies have reported widespread existence of RNA binding proteins that work as m^6^A *trans* factors to mediate the cell-specific m^6^A methylation through recruiting m^6^A methyltransferase complex (MTC) to the loci with the RBP binding^51^. These trans factors may bind to specific RNA isoforms and facilitate the m^6^A deposition on them. Additionally, these m^6^A trans factors include a number of canonical RNA splicing factors, such as SRSF7, RBFOX2, and TARBP2. SRSF7 can interact and colocalize with METTL3, METT14, and WTAP to facilitate the deposition of m^6^A near its binding sites^52^; while RBFOX2 can recruit RBM15, an MTC component, to chromatin RNAs^53^. Of note, TARBP2 can recruit MTC and promote the m^6^A methylation on TARBP2 bound transcripts^54^, the m^6^A will subsequently result in intron retention and RNA decay, which is consistent with our observation that specifically highly methylated isoforms are overrepresented with the intron retained isoforms. It is possible that the splicing factors mediate the complicated cross talk between alternative splicing and m^6^A modification. Fortunately, we have provided a powerful tool m6Aiso to decipher the detailed mechanisms that mediate the isoform-specific m^6^A modification.

## METHODS

### Cell culture

HeLa and HEK293T cells were cultured with Dulbecco’s modified Eagle medium (DMEM, GIBCO, Carlsbad, CA, USA) with 10% fetal bovine serum (FBS). To establish the model of DART-seq, HEK293T cells were transfected with pcDNA3.1-APOBEC1-YTH-T2A-copGFP plasmid by using Lipofiter 3.0 (Hanbio) according to the manufacturer’s instructions. After transfection 48 hours, the copGFP-positive cells were isolated through cytometry. To establish the model of cancer cells undergoing EMT, HeLa cells cultured with Dulbecco’s modified Eagle medium (DMEM, GIBCO, Carlsbad, CA, USA) without 10% fetal bovine serum (FBS) for 12 hours. Then, 10 ng/ml TGF-β was added to DMEM medium for 72 hours to induce HeLa cells undergoing EMT.

### ONT direct RNA sequencing and data preprocess

Nanopore direct RNA sequencing was performed in accordance with the guidelines provided by Oxford Nanopore Technologies (Oxford, UK) using DRS kits (SQK-RNA002) and R9.4.1 flow cells. The ionic current data for each FAST5 file were subjected to base calling using Guppy v4.2.2 with high accuracy model. Only reads exceeding the quality threshold of 7 were selected for subsequent analyses.

Sequencing reads were then aligned to the GRCh38 Ensembl annotated transcripts (v91) using minimap2 (v2.17)^55^ with the following parameters: ‘minimap2-ax map-ont-k14-uf-secondary=no--MD)’. Samtools (v1.9) was utilized to filter out secondary and supplementary alignments and convert the aligned reads to the BAM format^56^. ESPRESSO^57^ (v1.4.0) was used to measure the expression of transcripts by counts. To align the read signals with the corresponding transcript references, the *eventalign* module of Nanopolish’s (v0.13.2) was employed with the ‘--scale-events’ and ‘--signal-index’ options^58^. After the process of resquiggling, the continuous ionic current measurements from each read were segmented into 5-mer events comprised of the mean, standard deviation, and dwelling time along the corresponding transcriptome coordinates.

### Endogenously labeling of m^6^A on single reads and generation of training data

For the ONT DRS reads in HEK293T cells transfected with APOBEC1-YTH and empty vector, we used Sam2Tsv pileup (v23c0a5c) to identify the C-to-U mutations and calculate the mutation rates. To identify the APOBEC1-YTH induced C-to-U mutations, we firstly filtered out the mutations in dbSNP (v151) as well as the previously reported C-to-U mutations that could be induced by APOBEC1 alone^31^.

We further removed the C-to-U mutations that could be observed in the control cells with empty vector as determined by requiring the mutation rate exceeding 0.2 out of at least 10 reads. The remaining sites with C-to-U mutations on at least 10 reads and C-to-U mutation rate exceeding 0.05 but below 0.9 were considered as the preliminary APOBEC1-YTH induced C-to-U mutations for further analyses. To identify the clustered C-to-U mutations, we initially mapped the C-to-U mutations distributed across various isoforms of a single gene to the transcript with the longest coding regions. Then, we used 100 nt sliding windows with sliding step of 50 nt along the transcript with longest coding regions for each gene to search for C-to-U mutations. After merging the windows with C-to-U mutations, all the C-to-U mutations sites in the merged windows containing at least 3 C-to-U mutation sites were considered as clustered C-to-U mutations and preserved for the downstream analysis. For each clustered C-to-U mutation on each single read, the nearest GLORI annotated m^6^A site in DRACH motif in HEK293T cells^10^ within a 100 nt distance from the mutation was initially determined as the methylated m^6^A site on this single read. These methylated m^6^A sites on single reads were determined as the modified set of m^6^A on single ONT DRS reads only if there was an absence of C-to-U mutation within a 9 nt distance from the corresponding m^6^A sites on the same reads.

To determine the unmodified set of m^6^A on single ONT DRS reads of HEK293T cells transfected with APOBEC1-YTH, we first determined the unmodified 5-mers of DRACH by excluding the sites within a distance of 20 nt from the known SNPs in dbSNP (v151) and the known single-nucleotide m^6^A sites. Specifically, the previously determined single-nucleotide m^6^A sites in HEK293T cells by NGS-based experimental methods including GLORI^10^, miCLIP^33^, m6ACE-seq^34^, m6A-SAC-seq^9^ were excluded; then the peak regions of m6A-seq^35^ data in HEK293T cells determined as previously described^59^ were also excluded.

To generate an independent testing positive set of methylated m^6^A on single reads from the ONT DRS reads of HEK293T cells transfected with empty vector, we took the m^6^A sites on all the individual reads for the m^6^A sites with GLORI determined methylation levels exceeding 0.95 in HEK293T cells^10^.

The mean, standard deviation and dwell time of the ONT electric current signals at the 5-mers of DRACH motif as well as the 5-mers right upstream and downstream of the DRACH motif for the above generated modified set, unmodified set, and independent testing set were obtained for the downstream machine learning to develop m6Aiso.

### m6Aiso model and its learning parameters

The overall framework of m6Aiso is depicted in Supplementary Fig. 2. In brief, m6Aiso is a modified deep residual neural network model (ResNet) with the input of RNA sequences and their local-signal features to predict the states of sequence modification. In m6Aiso (Supplementary Fig.2), the 1*N-dimension input sequences were first transformed to 4*N-dimension matrix data using one-hot encoding method; then, a convolutional neural network (CNN) layer was designed to learn the high-level sequence features by using multiple convolution filters (4*5*2) sliding on the whole sequences with stride 1. After that, we concatenated the output of the learned high-level features of local sequences in each filter with their corresponding signal features and then inputted the combined feature matrix (5*N-4) to a 6-layer ResNet unit, which stacked by three ResNet blocks. The output of the last ResNet block was connected to a global maximum pooling layer and then connected to a two-layer fully-connected neural network with ReLU and sigmoid activation functions, respectively. In the first CNN layer, which accepts the input sequences, we used 2 convolution filters with size of 4*5; in the first ResNet block, we used 32 filters with size of (3ξ3) in all CNN layers; we used 128 filters with size of (3ξ3) in all CNN layers and 32 filters with size of (3ξ3) in all CNN layers in the second and third ResNet blocks, respectively. In each ResNet block, the dropout strategy was used in the forward and backward propagation between CNN layers and connected to a maximum pooling layer after each CNN layer. The first fully-connected layer was set to 16-way and the second fully-connected layer was set to 1-way to output the positive prediction. As a result, m6Aiso outputs the predicted modification probability at each DRACH sites. A read is deemed modified if its predicted modification probability surpasses a threshold of *P* = 0.9. Conversely, a site on an individual read with predicted modification probabilities < 0.1 are classified as unmodified, while the remainder are discarded.

### Model training and evaluation

By considering the situation that there are some false positive sequence samples in our training positive samples, we used a semi-supervised leaning strategy to train a better m6Aiso model. In detail, there are pseudo-labels for some positive samples, while all labels of negative samples are real in our training sequence data. Meanwhile, the number of positive samples versus the number of negative samples are extremely unbalance (1:25). It is hard to train a good prediction model if we use those data to train m6Aiso directly. To overcome the issues of pseudo-labels in positive samples and data unbalance in training data, we designed a semi-supervised learning strategy to train the m6Aiso model. Specifically, we first randomly selected a part of negative samples and combined them with all positive samples to constitute a new training data set with relative balance samples (positive vs. negative with 1:2.5); then we used the selected data set to train an initial prediction m6Aiso model and used it to predict the pseudo-label of each positive sample in the training data. After that, we then discard the samples with modified probabilities < 0.5 from the positive sample set and updated the training data to re-train the m6Aiso model. This update and re-training process will be repeated and terminated till the false positive rate (FPR) < 0.05 in the positive training data. Notably, although we used the trained model to re-predict the positive samples in the training data, this was different from the standard self-training pipeline in semi-supervised learning and may not be useless in most cases; while, as our training data skews to negative samples, the fitted model will tend to predict false positive samples with negative pseudo-labels. This will help us to obtain cleaner positive samples in training data gradually and fit more conservative model. To train the m6Aiso model, we used the cross-entropy loss function and Adma optimizer to optimize the weights in neural networks using learning rate of 0.001. In each m6Aiso model training, we set the batch size as 64, and the epoch as 50.

### Comparison between m6Aiso and NGS-based experimental methods

Based on the m6Aiso determined m^6^A sites on single reads, the genomic sites with at least 20 m^6^A modified reads were identified as m6Aiso determined m^6^A sites at gene level. The m^6^A sites identified by miCLIP^33^, m6A-SAC-seq^9^, and m6ACE-seq^34^ were obtained as previously provided. In addition, in order to evaluate the lowly methylated m^6^A sites, we also included the GLORI identified m^6^A sites with levels < 0.1 in the m^6^A level comparisons by re-executing the original bioinformatics pipeline of GLORI^10^ but without removing the sites with levels < 0.1.

### Comparison of m6Aiso with ONT-based computational tools

To evaluate the effectiveness of m6Aiso in mapping m^6^A modifications, we conducted a comparative analysis with established computational tools capable of detecting m^6^A at the read level and supporting methylation rates calculations. We ran m6Anet v2.0.0^29^ with default parameter, and only the DRACH sites with at least 20 reads were considered. For DENA v1.0^28^, we required at least 20 reads for the m^6^A sites and preserved the sites with modification level below 0.1. For nanom6A v2.0^27^, reads with a modification probability exceeding 0.8 were considered as modified in this study.

### Performance of m6Aiso on the synthetic Curlcakes dataset

To assess the capacity of m6Aiso to accurately distinguish between modified and unmodified reads, we performed inference using the synthetic Curlcakes dataset^21^. We combined all reads from two replicates of the modified and unmodified samples, respectively; and we excluded all 5-mer motifs that contained more than one adenosine. Thus, only reads from 43 sites within four DRACH motifs (GGACC, GGACU, UGACC, and UGACU) were preserved for subsequent analyses.

### Calculation of m^6^A methylation and expression levels for isoforms

For m6Aiso and m6Anet determined m^6^A sites at gene level, we calculated the m^6^A level of these sites on each isoform according to the aligned isoforms of the long reads that carry the m^6^A sites. At least 10 reads covering the m^6^A sites on the same isoforms were required to determine the methylation levels of the m^6^A sites on specific isoforms. Based on the m^6^A sites on each isoform, the m^6^A level of the isoform was calculated as the sum of all the modified reads at all m^6^A sites on the isoform divided by the sum of the read coverages of all those sites. Meanwhile, The C-to-U mutation level of each isoform was determined as the sum of the mutated read counts of all C-to-U mutation sites on the isoform divided by the sum of the read coverages of all those mutation sites.

### Analysis of wild-type and *METTL3*-KO samples

To assess the precision of m6Aiso, we obtained ONT-DRS data for wild-type and corresponding *METTL3*-KO in HEK293T and mESCs from the Singapore Nanopore-Expression project (SG-NEx project)^38^ and Zhong et al.^30^, respectively. The datasets underwent identical preprocessing steps as previously described, with the exception that the mESCs samples were aligned to the GRCm38 Ensembl annotated transcripts (v91). The m^6^A sites supported by at least 20 modified reads in WT or KO and covered by more than 50 reads in both conditions were used in the comparisons of m^6^A levels between WT and KO

### Co-occurrence analyses of m^6^A sites

Only the reads spanning at least 90% of the length of their corresponding mapped transcripts are used for the analyses. For each read and isoform, we calculated the transcriptomic distance between two adjacent m^6^A sites. For each gene we initially mapped the m^6^A sites to the isoform with the longest coding region and calculated the transcriptomic distance between each two adjacent sites.

For each pair of m^6^A sites in the same gene, we employed the parameter D’^39^, a matric commonly used to assess the linkage disequilibrium (LD) between two genetic variants within a population, to investigate the linkage of the pair of m^6^A sites across all the reads that covered the two sites using the following formula:

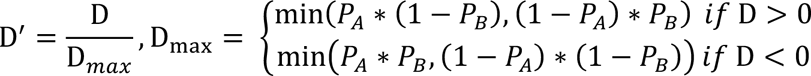

where D = P*_AB_* – P*_A_* P_B_, *P_A_* and *P_B_* represented the proportions of reads with modified A site and modified B site, respectively, while *P_AB_* denoted the proportion of reads with both modifications. A positive D’ value implies that the two modified sites tend to co-occur beyond random chance.

### Analysis of m^6^A sites on isoforms of single genes

For a specific m^6^A site on a particular isoform, we employed Fisher’s exact test to assess whether its m^6^A level was significantly differed from the combination of other isoforms from the same gene followed by Benjamini-Hochberg^60^ based false discovery rate (FDR) correction for multiple testing. The m^6^A sites exhibiting m^6^A level changes exceeding 0.1 and FDR less than 0.05 were considered as isoform-specific highly or lowly methylated m^6^A sites. To analyze the transcript type compositions of the isoform-specific highly and lowly methylated m^6^A sites, we excluded the genomic sites that exhibit highly methylated on one isoform while displaying lowly methylated on the other isoform.

To investigate the relationship between m^6^A and their proximity to exon junctions, we compared the m^6^A levels of identical sites on pairs of different isoforms. We exclusively used the sites that were more than 200 nt away from EJCs on one of the pairs of isoforms but less than 100 nt away from the EJC on the other isoforms. Fisher’s exact test was used to evaluate the difference in modification level of m^6^A sites, with FDRs calculated using Benjamini-Hochberg^60^ method. The m^6^A sites were considered as differentially methylated m^6^A sites if the absolute changes of m^6^A levels between two isoforms exceeding 0.1 and FDRs less than 0.05.

### Identification of dynamic m^6^A sites during EMT process

Cutadapt^61^ (v3.5) was used to remove the adapters from the Illumina sequencing reads. Then, the reads were aligned to human reference genome (GRCh38) using STAR^62^ (v2.7.4a). FeatureCounts^63^ (v2.0.1) was utilized to quantify the number of reads mapped to each gene. Genes with counts exceeding 10 in both control and TGF-β samples were retained for subsequent analysis. Differential expression analysis was conducted using DESeq2^64^. The genes with fold changes > 2, and *P*-adjust values < 0.01 were identified as the differentially expressed during EMT process. GO enrichment analysis was conducted using the R package Clusterprofile (v4.3.8)^65^.

For both replicates of the ONT DRS control and TGF-β samples, we preprocessed them as described above previously. The m6Aiso determined common m^6^A sites of the two replicates with identical loci on the genome in control or TGF-β samples were used in the downstream isoform analyses. To investigate isoform-specific dynamic changes of identical m^6^A sites, we compared the same m^6^A sites on the same isoforms between control and TGF-β samples. Only the m^6^A sites on isoforms covered at least 20 reads in all samples were used in the analyses.

Differentially methylated m^6^A sites at isoform level were determined as the sites with absolute changes in their modification levels between control and EMT cells exceeding 0.15. Genes containing dynamically modified sites were selected for GO enrichment analysis using clusterProfiler (v4.3.8)^65^. The fractions of expressed isoforms for each gene were calculated based on the ESPRESSO (v1.4.0)^57^ determined isoform expression.

## DATA AVAILABILITY

All data generated for this paper has been deposited in the NCBI Sequence Read Archive (SRA) database under accession number PRJNA1044456 ( reviewer accessible link: https://dataview.ncbi.nlm.nih.gov/object/PRJNA1044456?reviewer=egtn8d6qio41rs72mcu1ms2mko). The synthetic Curlcakes datasets are obtained the work of Liu et al.^21^ and are available through the NCBI Gene Expression Omnibus (GEO) database under accession number GSE124309. Published DRS datasets derived from WT and *Mettl3* KO mESCs samples are obtained the work of Zhong et al.^30^ and are available at the GEO database under accession number GSE195618. Published DRS datasets for WT and *METTL3* KO HEK293T samples are from the work of Pratanwanich et al.^25^ and are available through European Nucleotide Archive (ENA) database under accession number PRJEB40872.

## CODE AVAILABILITY

The source code for m6Aiso is available on GitHub at https://github.com/Jinkai-Wang-Lab-epitranscriptomics/m6Aiso.

## ACKNOWLEDGEMENTS

This work was supported by the National Natural Science Foundation of China (32270630, J.W.; 31970594, J.W.; 32300455, Y.C.; 62102173, Y.G.), China Postdoctoral Science Foundation (2023M734053, X.H.), the Guangdong Science and Technology Program (2022A1515110255, Y. C.), the Fundamental Research Funds for the Central Universities (23xkjc003, J.W.; 22qntd4810, Y.C.; 23ptpy90, Z.R.; 23ptpy87, X.H.)

## Supplementary Figures

**Supplementary Figure 1.**
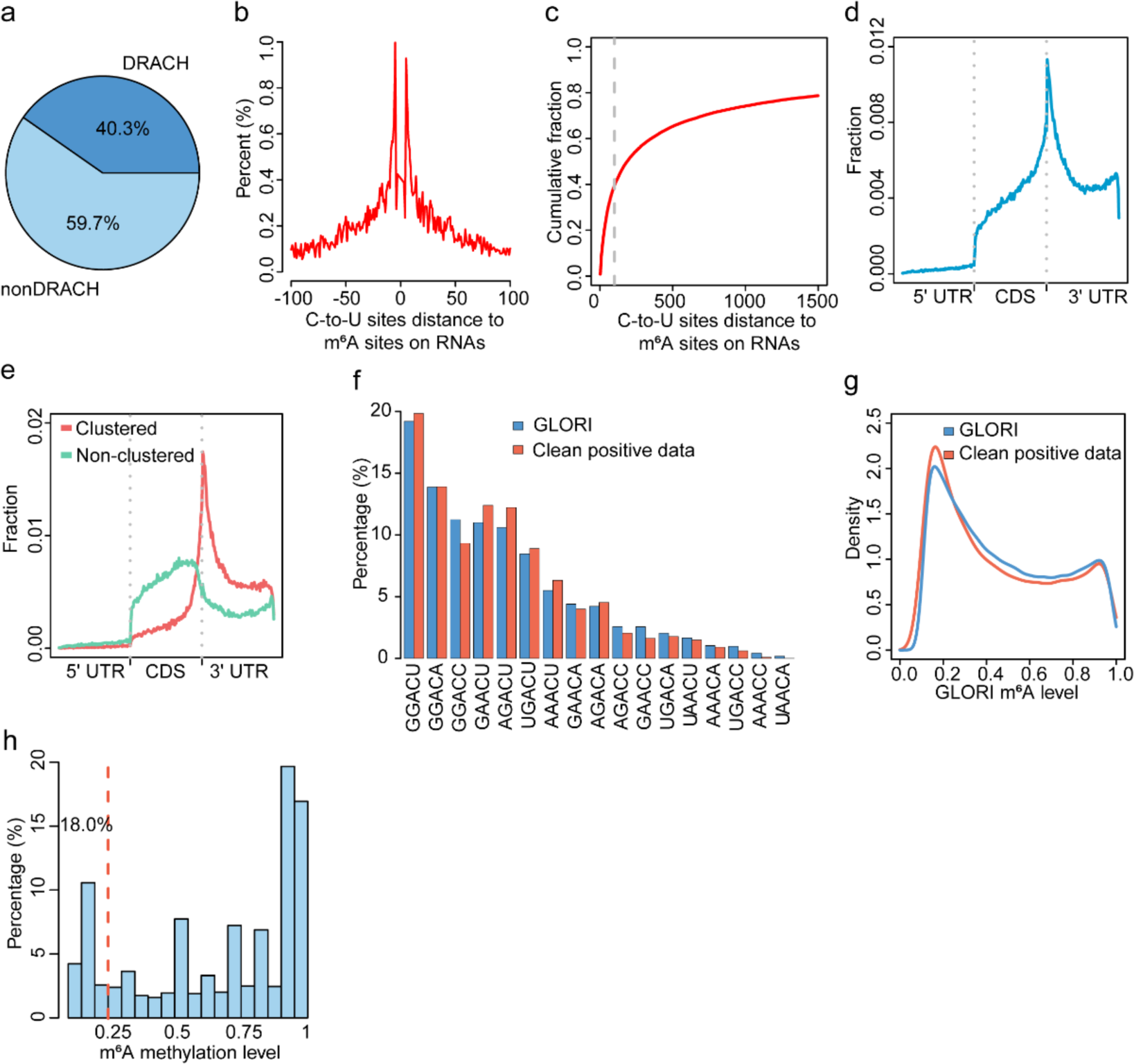
Characteristics of positive signals obtained from endogenously labeled individual DRS reads. **a**, Pie chart displaying the proportion of C-to-U mutations occurrence within and outside the DRACH motif. **b**, The distribution of the distances of C-to-U mutations relative to GLORI annotated m^6^A sites on RNAs. **c**, Plot of cumulative fraction of the distances between C-to-U mutated sites and GLORI annotated m^6^A sites on RNAs. **d**, Metagene plot illustrating the distribution of the C-to-U sites identified from DRS data. **e**, Metagene plot illustrating the distribution of the clustered and non-clustered C-to-U sites from DRS data. **f**, Comparison of the percentages of unique m^6^A sites in the clean positive training data with GLORI annotated m^6^A sites on the DRACH motifs. **g**, Density plot comparing the distributions of m^6^A methylation levels at sites from the clean positive data and GLORI. **h**, Histogram illustrating the percentages of m^6^A sites with varying methylation levels in the clean positive data.

**Supplementary Figure 2.**
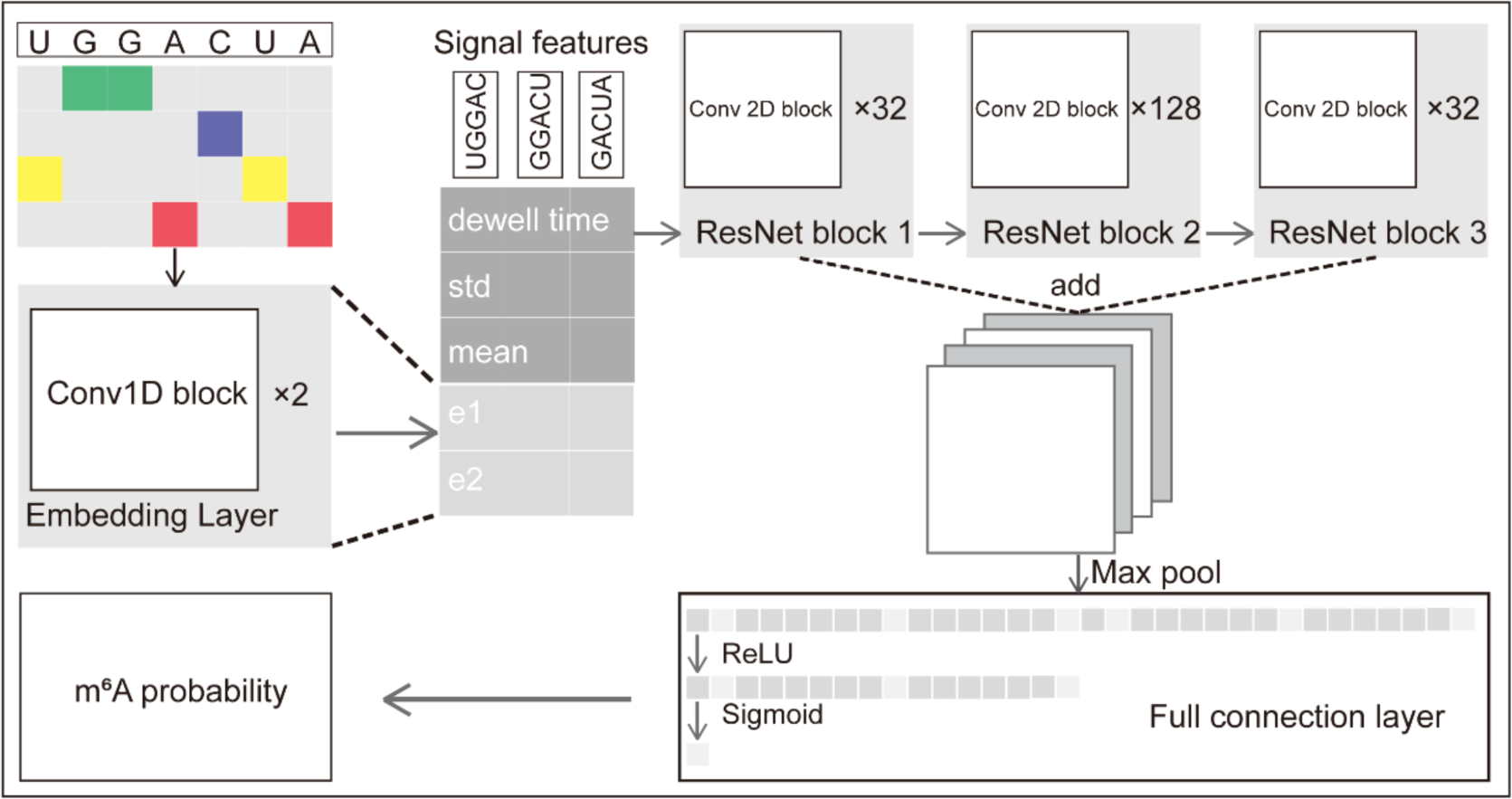
Detailed architecture of m6Aiso.

**Supplementary Figure 3.**
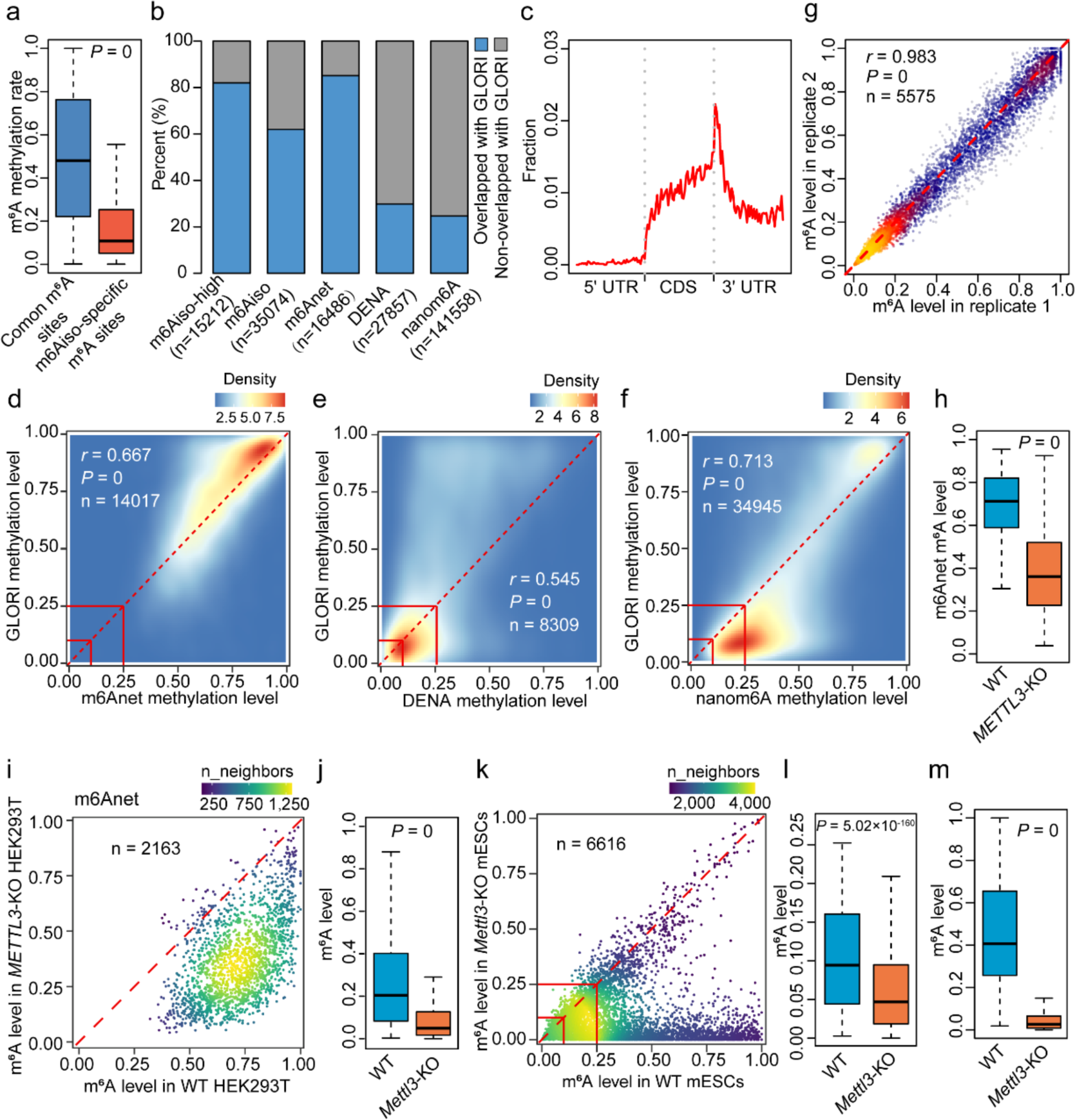
Performance comparison between m6Aiso and exiting computational tools. **a**, Box plot comparing the methylation levels between m^6^A sites specifically found by m6Aiso and the common m^6^A sites found by both m6Aiso and other methods. **b**, Percentages of m^6^A sites captured by the computationally methods that are validated by GLORI. “m6Aiso-high” denotes the m6Aiso determined m^6^A sites with methylation levels > 0.4. **c**, Metagene profile illustrating the distribution of the m6Aiso-specific modified sites. **d-f**, Correlation of the m^6^A levels estimated by m6Anet (**d**), DENA (**e**), nanom6A (**f**) and GLORI in HEK293T cells, respectively. The m^6^A levels of 0.1 and 0.25 are indicated by red lines. **g**, Correlation of the methylation levels for m^6^A sites detected by m6Aiso between the two WT HEK293T replicates. **h**, **i**, Box plot (**h**) and scatter plot (**i**) illustrating a decrease in methylation levels for m^6^A sites detected by m6Anet in *METTL3*-KO HEK293T cells compared to WT HEK293T cells. *P* value was calculated by two-tailed Wilcoxon rank-sum test. **j**, **k**, Box plot (**j**) and scatter plot (**k**) demonstrating a remarkable reduction of m^6^A levels in *Mettl3*-KO mESCs compared to WT mESCs for all the m6Aiso determined m^6^A sites. *P* value was calculated by two-tailed Wilcoxon rank-sum test. **l**, Box plot illustrating a decrease in methylation levels for the m6Aiso determined lowly methylated m^6^A sites (level < 0.25) in *Mettl3*-KO mESCs compared to WT mESCs. *P* value was calculated by two-tailed Wilcoxon rank-sum test. **m**, Box plot depicting a remarkable reduction of methylation levels in *Mettl3*-KO mESCs compared to WT mESCs for the GLORI-annotated m^6^A sites. *P* value was calculated by two-tailed Wilcoxon rank-sum test. For boxplots: center line, median; box limits, upper and lower quartiles; whiskers, 1.5x interquartile range.

**Supplementary Figure 4.**
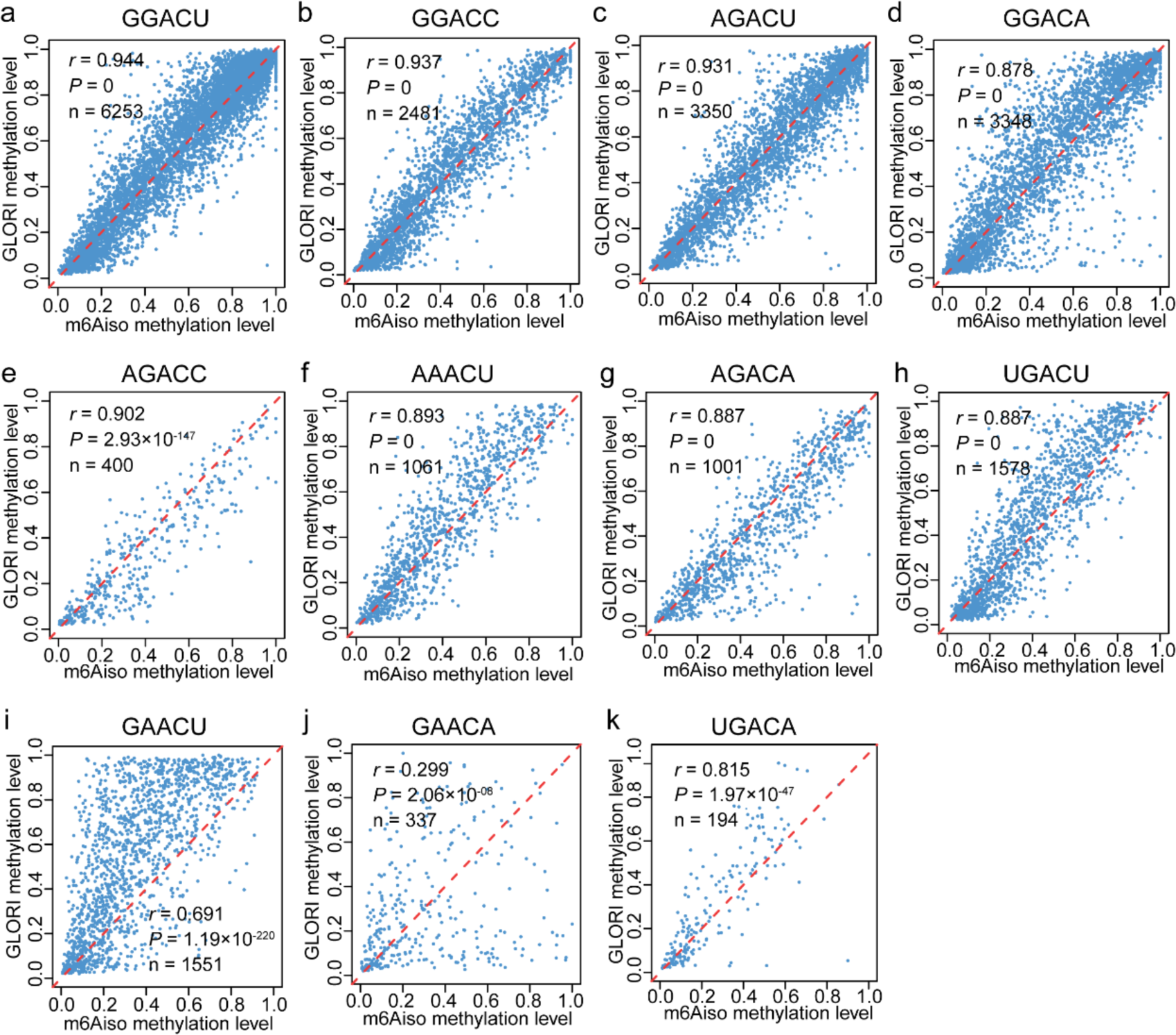
Correlation of the m^6^A levels estimated by m6Aiso and GLORI in HEK293T cells. **a-k**, Scatter plot illustrating the correlation of the m^6^A levels in HEK293T cells estimated by the method GLORI (y-axis) and m6Aiso (x-axis) in different 5-mers.

**Supplementary Figure 5.**
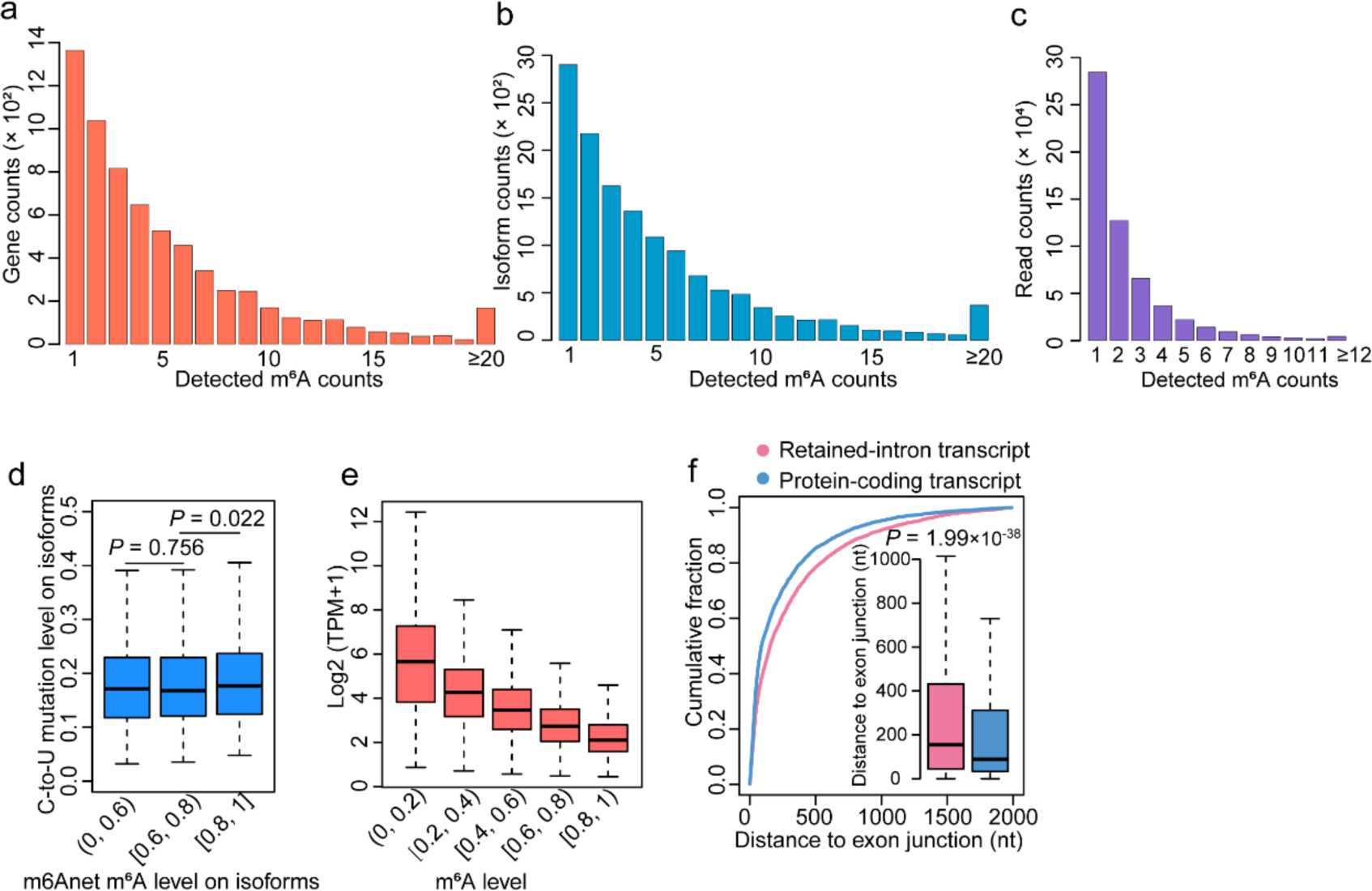
m6Aiso detects m^6^A sites at various levels. **a-c**, Frequency of genes (**a**), isoforms (**b**) and reads (**c**) containing different number of m^6^A sites. **d**, Correlation of isoform m^6^A levels determined by m6Anet and APOBEC1-YTH induced C-to-U mutations. *P* values were calculated by two-tailed Wilcoxon test. **e**, Correlation of isoform m^6^A level determined by m6Aiso and expression abundance. **f**, Plot comparing the cumulative fractions of EJDs for identical m^6^A sites on retained-intron isoforms and their corresponding protein-coding isoforms. *P* value was calculated by two-tailed Wilcoxon test. For boxplots: center line, median; box limits, upper and lower quartiles; whiskers, 1.5x interquartile range.

**Supplementary Figure 6.**
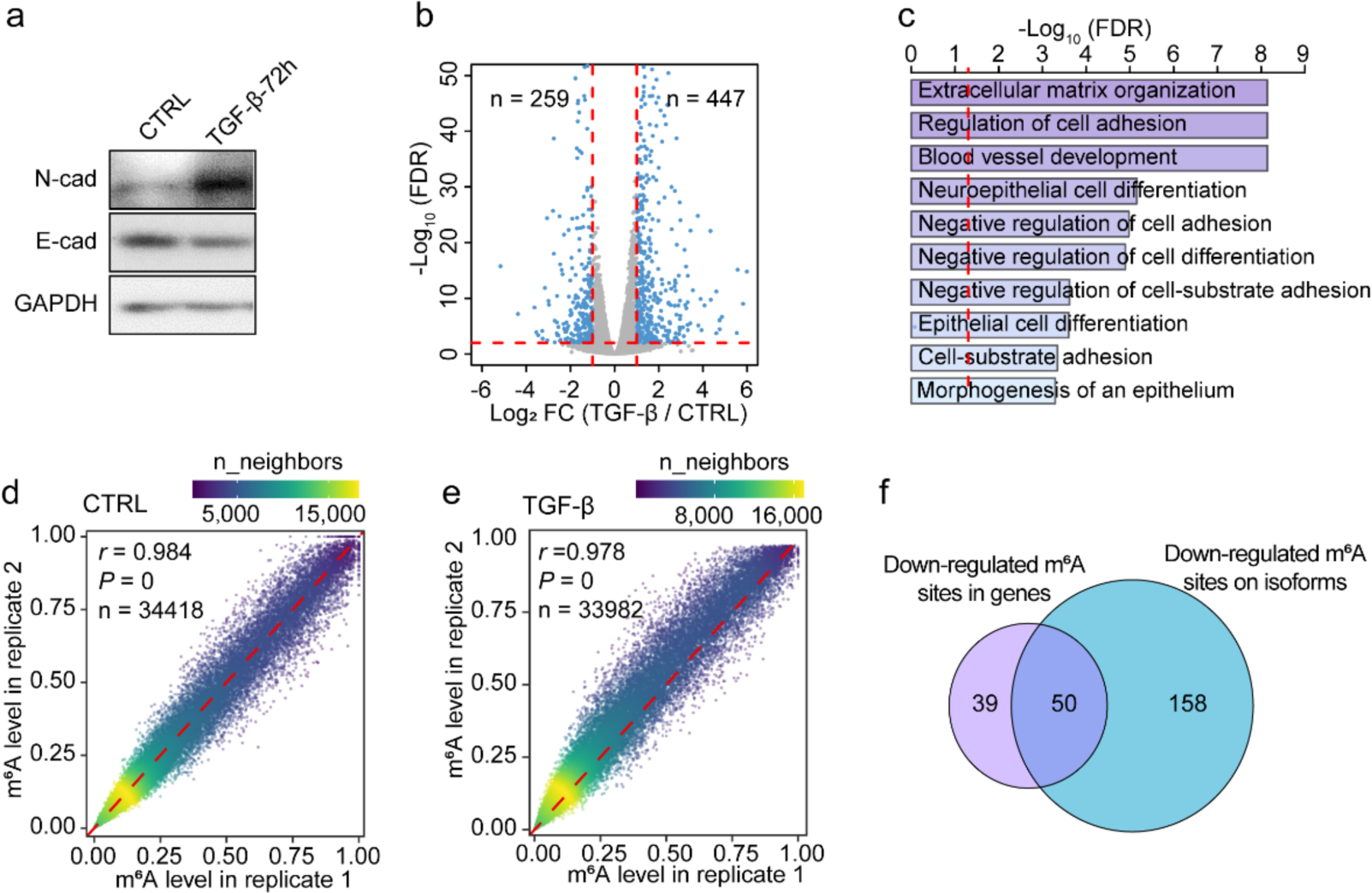
Establishment of an EMT model in HeLa cells by TGF-β treatment. **a**, Protein expression of N-cad and E-cad in control and TGFB-β induced cells were measured by western blot analysis. CTRL, control. **b**, Volcano plot depicting differential gene expression during TGF-β induced EMT. **c**, GO enrichment for the up-regulated genes undergoing EMT process. **d**, **e**, Correlation of the methylation levels for m^6^A sites detected by m6Aiso in two replicates of control (**d**) and TGF-β induced cells (**e**), respectively. **f**, Venn diagram comparing the down-regulated m^6^A sites at the gene and isoform level.

